## Supplementary Material for "Decoding Living Systems: Reassessing Crop Model Frontiers via Biological Dynamics and Optimized Phenotype"

Supplementary Material
Edgar S. Correa

#### Sensitivity Analysis

Sensitivity analysis was performed using 20 independent replications of the Morris method with randomized parameter sequences. (Table S1) presents complete RSI statistics (mean ± SD) for all parameter-output combinations.

**Key findings:**

- Mean 95% CI width = 0.04, confirming robust parameter rankings
- Phenological parameters (P1, PHINT) dominate developmental timing and biomass accumulation
- Reproductive parameters (G3, G2) control yield components but exhibit negligible influence on biomass (RSI $< 0.07$), confirming source-sink independence in CERES-Rice
- Thermal stress parameters (TCLDP, TCLDF) show negligible sensitivity under studied conditions

**Parameter definitions:**

- P1: Vegetative phase thermal time (GDD)
- P2O: Critical photoperiod (hours)
- P2R: Photoperiod delay coefficient
- P5: Grain filling duration (GDD)
- G1: Spikelet number coefficient
- G2: Potential grain weight (g)
- G3: Tillering coefficient
- PHINT: Phyllochron interval (GDD)
- THOT: Heat stress threshold (°C)
- TCLDP: Cold stress threshold - panicle (°C)
- TCLDF: Cold stress threshold - flowering (°C)

#### Operators in Genetic Algorithms

To maximize crop yield, grain conversion efficiency, and water use efficiency, a genetic algorithm (GA) is implemented to iteratively optimize the model parameters. The fitness function guides the processes of selection, recombination, and mutation, ensuring convergence toward parameter sets that balance agronomic performance and genotypic diversity.

1. **Selection Operator**

   Roulette Wheel Selection, or fitness proportional selection, is a probabilistic method in genetic algorithms. It starts by calculating the fitness of each individual in the population, which reflects how well they solve the optimization problem. The total fitness $F_{\text{total}}$ of the population is calculated by adding the fitness values of all individuals $f_i$. The selection probability $P_i$ for each individual $i$ is then computed by dividing the fitness of the individual $f_i$ by the total fitness $F_{\text{total}}$, as shown in Eq 1.

$$P_i = \frac{f_i}{\sum_{i=1}^{n} f_i} \tag{1}$$

Next, a random number $r \in [0, 1]$ is generated, and the cumulative selection probability $C_i$ is computed by summing the selection probabilities up to individual $P_i$. The individual $P_i$ is selected if the random number $r$ lies within the cumulative probability interval $[C_{i-1}, C_i)$, as defined in Eq 2.

$$C_{i-1} \leq r < C_i, \quad \text{where } C_0 = 0 \text{ and } C_n = 1 \tag{2}$$

This process ensures that individuals with higher fitness have a greater chance of selection, while still allowing for genetic diversity by giving less fit individuals a small chance to be selected.

2. **Recombination and mutation Operators**

   Arithmetic Crossover is a recombination operator. The primary objective of this operator is to combine two parent solutions to produce offspring solutions by generating new combinations of their values. Unlike traditional crossover methods that exchange segments or genes, arithmetic crossover blends the parent solutions through a weighted average. Given two parent solutions, $P_1 = \{P_1^1, P_1^2, \ldots, P_1^n\}$ and $P_2 = \{P_2^1, P_2^2, \ldots, P_2^n\}$, where each $P_1^i$ and $P_2^i$ represent the values of the corresponding genes in the solution vectors, arithmetic crossover produces a new offspring $O$ based on a linear combination of the parents' gene values. The new gene values in the offspring are calculated as expressed in Eq 3.

$$O^i = \alpha P_1^i + (1 - \alpha) P_2^i, \quad \forall i \in \{1, 2, \ldots, n\} \tag{3}$$

   where $\alpha$ is a random scalar weight factor in the interval [0, 1] that controls the influence of each parent on the offspring. Finally, a random mutation is applied, prioritizing the crop parameters with the highest sensitivity, based on the sensitivity analysis.

### Supplementary Note: Vapor Pressure Deficit (VPD) Calculation

Vapor pressure deficit was calculated using the Tetens equation [2, 4]:

$$\text{VPD} = e_s - e_a \tag{4}$$

where $e_s$ is the saturation vapor pressure and $e_a$ is the actual vapor pressure, calculated as:

$$e_s = 0.6108 \times \exp\left(\frac{17.27 \times T}{T + 237.3}\right) \tag{5}$$

$$e_a = e_s \times \frac{RH}{100} \tag{6}$$

with $T$ = mean temperature (°C) and $RH$ = relative humidity (%).

**Climate Type 1 (Southern humid zone):**

- $T = 26.39$°C, $RH = 80.33\%$
- $e_s = 0.6108 \times \exp(17.27 \times 26.39 / 263.69) = 3.43$ kPa
- $e_a = 3.43 \times 0.8033 = 2.76$ kPa
- **VPD = 0.67 kPa**

**Climate Type 2 (Northern drought-prone zone):**

- $T = 27.78$°C, $RH = 67.98\%$
- $e_s = 0.6108 \times \exp(17.27 \times 27.78 / 265.08) = 3.71$ kPa
- $e_a = 3.71 \times 0.6798 = 2.52$ kPa
- **VPD = 1.19 kPa**

The 77% higher VPD in northern environments (1.19 vs 0.67 kPa) indicates substantially greater atmospheric water demand, intensifying drought stress during critical reproductive stages. VPD values exceeding 1.0 kPa are generally associated with significant transpiration-driven water stress in rice [1, 3].

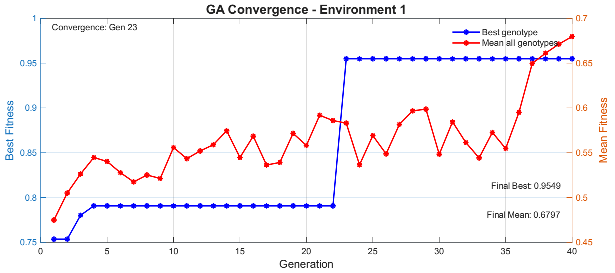


**(a)** Environment 1: high water retention (SDUL = 0.30), high precipitation (30% area)

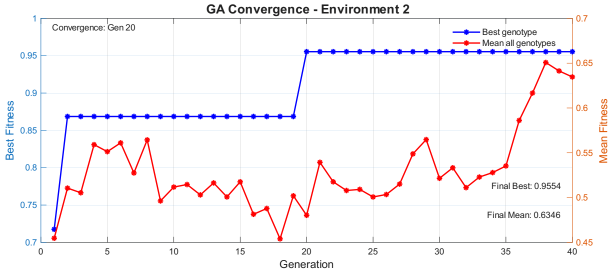


**(b)** Environment 2: medium retention (SDUL = 0.28), low precipitation (18% area)

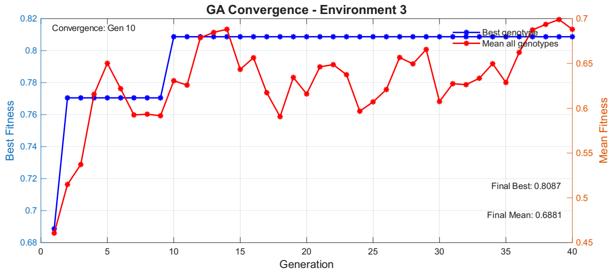


FigS1c.tif

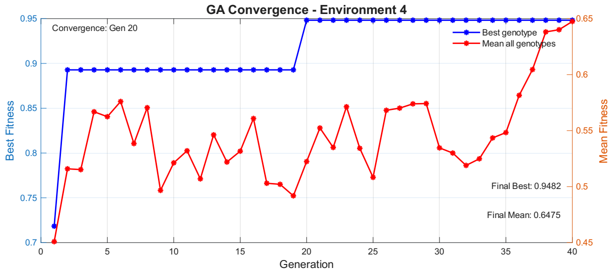


**(c)** Environment 4: low retention (SDUL = 0.23), low precipitation (20% area)

**Fig S1.** Genetic algorithm convergence across four environments. Blue line: best individual fitness per generation. Red line: population mean fitness. Convergence to 95% of maximum fitness occurred at generations 23, 20, 10, and 20 for Environments 1–4, respectively. Environments 2 and 4 yielded identical optimal genetic coefficients despite differing soil water retention.
