## Supplementary material for "Decoding Living Systems: Reassessing Crop Model Frontiers via Biological Dynamics and Optimized Phenotype": TableS1

**Table S1.** Relative Sensitivity Index (RSI) from 20 Morris method replications. Bold = highest sensitivity per output.

| **Coefficient** | **Grain Yield** | **Biomass** | **Anthesis** | **Maturity** | **N° Grains** | **N° Tillers** |
|---|---|---|---|---|---|---|
| P1 | 0.46 ± 0.04 | 0.55 ± 0.04 | **0.93 ± 0.02** | 0.57 ± 0.19 | 0.46 ± 0.04 | 0.88 ± 0.02 |
| P2O | 0.42 ± 0.04 | 0.45 ± 0.04 | 0.79 ± 0.12 | **0.64 ± 0.08** | 0.43 ± 0.04 | 0.76 ± 0.11 |
| P2R | 0.37 ± 0.03 | 0.39 ± 0.03 | 0.00 ± 0.00 | 0.50 ± 0.05 | 0.37 ± 0.03 | 0.05 ± 0.03 |
| P5 | 0.42 ± 0.04 | 0.42 ± 0.04 | 0.70 ± 0.05 | 0.35 ± 0.06 | 0.42 ± 0.04 | 0.66 ± 0.05 |
| G1 | 0.45 ± 0.04 | 0.06 ± 0.03 | 0.00 ± 0.00 | 0.00 ± 0.00 | 0.46 ± 0.04 | 0.00 ± 0.00 |
| G2 | 0.44 ± 0.04 | 0.05 ± 0.02 | 0.00 ± 0.00 | 0.00 ± 0.00 | **0.64 ± 0.03** | 0.00 ± 0.00 |
| G3 | **0.59 ± 0.04** | 0.48 ± 0.05 | 0.00 ± 0.00 | 0.00 ± 0.00 | 0.59 ± 0.04 | **0.93 ± 0.01** |
| PHINT | 0.42 ± 0.02 | **0.70 ± 0.02** | 0.86 ± 0.02 | 0.47 ± 0.05 | 0.42 ± 0.02 | 0.84 ± 0.03 |
| THOT | 0.45 ± 0.06 | 0.05 ± 0.02 | 0.00 ± 0.00 | 0.00 ± 0.00 | 0.45 ± 0.06 | 0.00 ± 0.00 |
| TCLDP | 0.00 ± 0.00 | 0.00 ± 0.00 | 0.00 ± 0.00 | 0.00 ± 0.00 | 0.00 ± 0.00 | 0.00 ± 0.00 |
| TCLDF | 0.00 ± 0.00 | 0.00 ± 0.00 | 0.00 ± 0.00 | 0.00 ± 0.00 | 0.00 ± 0.00 | 0.00 ± 0.00 |
