## Supplementary material for "Decoding Living Systems: Reassessing Crop Model Frontiers via Biological Dynamics and Optimized Phenotype": TableS2

**Table S2.** Pearson correlation ($R^2$) between HI-WUE index and phenotypic outputs across four environments. Statistical significance: *** $p < 0.001$, ** $p < 0.01$, * $p < 0.05$, ns $p \geq 0.05$. Correlations with $R^2 < 0.10$ are considered practically negligible despite statistical significance due to large sample sizes ($n > 1{,}300$ per environment).

| **Env** | **Metric** | **GY** | **Biomass** | **HI** | **N°Grains** | **LAI** | **WUE** | **Root** | **Anthesis** | **Maturity** |
|---|---|---|---|---|---|---|---|---|---|---|
| 1 | $R^2$ | 0.78 | 0.04 | **0.88** | 0.39 | 0.01 | **0.86** | 0.02 | 0.00 | 0.01 |
| | p-value | *** | *** | *** | *** | ns | *** | ** | ns | * |
| 2 | $R^2$ | 0.86 | 0.03 | **0.97** | 0.56 | 0.05 | **0.93** | 0.02 | 0.06 | 0.08 |
| | p-value | *** | *** | *** | *** | *** | *** | ** | *** | *** |
| 3 | $R^2$ | 0.78 | 0.00 | **0.95** | 0.60 | 0.00 | **0.88** | 0.00 | 0.00 | 0.00 |
| | p-value | *** | ns | *** | *** | ns | *** | ns | ns | ns |
| 4 | $R^2$ | 0.85 | 0.02 | **0.97** | 0.50 | 0.03 | **0.93** | 0.01 | 0.04 | 0.05 |
| | p-value | *** | ** | *** | *** | *** | *** | * | *** | *** |
