## Supplementary material for "Decoding Living Systems: Reassessing Crop Model Frontiers via Biological Dynamics and Optimized Phenotype": TableS3

**Table S3.** Optimal genetic coefficients and predicted crop performance for each environment. Env 1: high water retention, high precipitation (30% area); Env 2: medium retention, low precipitation (18%); Env 3: medium retention, high precipitation (21%); Env 4: low retention, low precipitation (20%). Values represent the best individual identified across 40 generations of genetic algorithm optimization.

| **Parameter** | **Range** | **Env 1** | **Env 2** | **Env 3** | **Env 4** | **Description** |
|---|---|---|---|---|---|---|
| *Genetic coefficients* | | | | | | |
| P1 (GDD) | 150–800 | 526.6 | 605.0 | 538.1 | 605.0 | Vegetative phase thermal time |
| P5 (GDD) | 150–850 | 372.1 | 207.0 | 247.8 | 207.0 | Grain filling duration |
| P2R (GDD) | 5–300 | 237.4 | 112.8 | 204.6 | 112.8 | Photoperiod sensitivity |
| PHINT (GDD) | 55–90 | 73.9 | 72.0 | 74.1 | 72.0 | Phyllochron interval |
| P2O (h) | 11–13 | 11.7 | 11.7 | 12.1 | 11.7 | Critical photoperiod |
| G1 (#/g) | 38–540 | 64.0 | 61.0 | 61.0 | 61.0 | Spikelet number coefficient |
| G2 (g) | 0.015–0.030 | 0.025 | 0.025 | 0.027 | 0.025 | Single grain weight |
| G3 | 0.7–1.97 | 0.85 | 0.97 | 0.95 | 0.97 | Tillering coefficient |
| *Predicted performance* | | | | | | |
| Grain yield (kg/ha) | | **4,837** | 4,014 | 4,213 | 3,743 | |
| Biomass (kg/ha) | | 8,895 | 7,200 | 7,214 | 6,802 | |
| HI | | 0.54 | 0.56 | **0.58** | 0.55 | |
| WUE ($kg\ ha^{-1} mm^{-1}$) | | **6.17** | 5.97 | 5.84 | 5.97 | |
| Grain number (/$m^2$) | | 19,350 | 16,056 | 15,602 | 14,971 | |
| Tiller number (/$m^2$) | | 956 | 1,253 | 906 | 1,242 | |
| Anthesis (days) | | 87 | 80 | 80 | 80 | |
| Maturity (days) | | 116 | 101 | 103 | 101 | |
| LAI ($m^2/m^2$) | | 3.73 | 3.23 | 2.97 | 3.03 | |
| *Optimization metrics* | | | | | | |
| Best fitness (HI-WUE) | | 0.955 | 0.955 | **0.973** | 0.948 | |
| Convergence (gen) | | 23 | 20 | 10 | 20 | Generation at 95% max fitness |
